# High-Throughput Evaluation of Cryoprotective Agents for Mixture Effects That Reduce Toxicity

**DOI:** 10.1101/2025.05.02.651925

**Authors:** Nima Ahmadkhani, Cameron Sugden, James D. Benson, Ali Eroglu, Adam Z. Higgins

## Abstract

Vitrification is a promising approach for cryopreserving complex biological structures such as organs. However, to prevent ice formation, high concentrations of cell-permeable cryoprotective agents (CPAs) are required, which can be highly toxic. The current reliance on a small number of CPAs limits optimization of low-toxicity compositions for vitrification. To address this, there is growing interest in uncovering novel chemicals with analogous protective qualities. This may not only enhance vitrification efficacy but also mitigate toxic effects. In the current study, we employed a high throughput method to assess the toxicity of 21 compounds at room temperature, both individually and in binary combinations. Our analysis revealed that toxicity increases with both exposure duration and concentration, and that several CPA combinations result in reduced overall toxicity. Notably, among all tested mixtures, four binary combinations—formamide/glycerol, dimethyl sulfoxide/1,3-propanediol, 1,2-propanediol/diethylene glycol, and 1,3-propanediol/diethylene glycol—produced a statistically significant decrease in toxicity, resulting in significantly higher viability for the 6 mol/kg mixture than both corresponding 6 mol/kg single CPA solutions. The high-throughput approach presented here will aid in building a comprehensive CPA toxicity database, which will improve our understanding of toxicity mechanisms and support the development of predictive models for identifying novel CPA mixtures with low toxicity.

## 1. Introduction

Organ transplantation is a life-saving procedure, but the field encounters substantial obstacles, such as a shortage of organs, limited preservation times, and a risk for organ failure resulting from inadequate preservation techniques [19,25]. The effectiveness of organ transplantation relies heavily on the proper preservation of organs from the moment they are collected until they are implanted [8,9]. Conventional preservation techniques, typically involving hypothermic storage, offer only a limited window for organ viability. This often results in suboptimal outcomes and increased risks of transplant failure [10,18].

To tackle these difficulties, cryopreservation has arisen as a possible remedy. Cryopreservation involves cooling biological materials to very low temperatures, effectively halting metabolic and chemical processes [8,32]. This technology has the potential to significantly prolong the viability of organs and tissues beyond the constraints of other preservation procedures. Cryopreservation could significantly impact the field of organ transplantation by allowing for more flexible scheduling of transplants, improving donor-recipient matching, and enhancing overall transplant success rates [9,22].

Vitrification has shown significant potential for organ cryopreservation [22,35]. Vitrification is a method that involves fast cooling and warming and high concentrations of cryoprotective agents (CPAs) to transform biological materials into a glassy, non-crystalline state. This technique is different from standard freezing, which might damage cells by forming ice crystals. Vitrification prevents ice formation and growth, therefore reducing cellular damage and maintaining structural integrity [5,32,35]. Vitrification has been successfully applied to a range of biological specimens, from embryos to complex tissues, demonstrating its potential for improving organ preservation [27,35]. In a recent study, Han et al. achieved reproducible cryopreservation of rat kidneys via vitrification. They demonstrated that the kidneys remained functional after being transplanted into rats [22]. Other research studies documented the survival of vitrified rabbit kidneys following transplantation [16,36]. Both cases showed signs of kidney damage, highlighting the need for improved cryopreservation methods. Moreover, a major obstacle in translating these results to human organs is their larger size, which results in slower rates of cooling and warming. Thus, a greater concentration of CPAs is required to prevent ice formation in human organs [21].

As a result, there is significant interest in developing CPA formulations that are less toxic at high concentrations. A common strategy is to use multi-CPA solutions because mixtures have been shown to reduce overall toxicity compared to single-CPA solutions [4,11,13,33,35]. Toxicity reduction in mixtures has been attributed to two main mechanisms: mutual dilution and toxicity neutralization. Mutual dilution describes how each CPA in a mixture lowers the concentration of the other CPAs. Because CPA toxicity increases with concentration, mutual dilution can lead to reduced toxicity for the mixture compared to the equivalent concentration of the constituent CPAs [13,33,35]. Toxicity neutralization is defined as the reduction or elimination of one CPA’s toxicity through the addition of a second CPA [11]. Previous studies to explore toxicity reduction in CPA mixtures have been limited in scope, highlighting the need for further studies to identify promising CPA combinations.

In our previous study, we conducted a comprehensive evaluation of the toxicity of five frequently used CPAs – formamide, 1,2-propanediol (aka propylene glycol), ethylene glycol, dimethyl sulfoxide, and glycerol – along with combinations of these CPAs in ternary and binary forms [35]. We examined the impact of concentration and duration of exposure to these CPAs at room temperature on bovine pulmonary artery endothelial cells (BPAECs), which were used as a model system. A Hamilton Microlab STARlet system was used to automate liquid handling operations, improving both the accuracy and throughput of the experiments compared to manual approaches. Additionally, the automation allowed for the randomization of CPA treatments in the 96-well plate. Our results confirmed neutralization of formamide toxicity by dimethyl sulfoxide, as previously observed by Fahy [11], and revealed a new case of toxicity neutralization in mixtures containing formamide and glycerol. This underscores the potential for using automated liquid handling technology to streamline identification of promising CPA combinations.

Prior research on cryopreservation of tissues and organs have focused on a limited number of CPAs [7,26]. Effective CPAs have many common traits, including low toxicity, the ability to penetrate cell membranes, and the prevention of ice formation in aqueous solutions [23,35]. There exists a multitude of chemicals that have not yet undergone testing but have the potential to be beneficial for cryopreservation [1,28,29].

To screen for new molecules that can serve as CPAs, we recently developed a new method for high throughput measurement of CPA permeability and toxicity. The method relies on volume-dependent changes in calcein fluorescence measured using an automated plate reader. This technique was used to investigate the permeability and toxicity of 27 chemicals at both 4 °C and room temperature at low concentrations (1 and 2 mol/kg) [1]. Of these 27 molecules, 21 had high permeability and low toxicity at room temperature.

In this investigation, we assessed the toxicity of these 21 molecules [1] at higher concentrations using the Hamilton high throughput approach [35]. The objective was to determine the toxicity of these CPAs, both individually and in binary mixtures, to identify compositions with low toxicity. This approach allowed us to assess a wide range of CPA combinations for their ability to reduce toxicity.

## 2. Materials and Methods

### Materials

The 21 CPAs used in this study were selected from the 27 CPAs screened by Ahmadkhani et al [1] (see Table 1). Other chemicals and reagents, including PrestoBlue, HEPES buffered saline, Dulbecco’s Modified Eagle Medium (DMEM), fetal bovine serum, and penicillin-streptomycin, were sourced as described by Warner et al [35].

**Table 1.** CPA exposure conditions tested

| CPAs tested at 3 mol/kg | CPAs tested at 6 mol/kg |
| --- | --- |
| 1,3-Dihydroxyacetone (DHA) | 1,3-Propanediol (PD) |
| 1,3-Propanediol (PD) | 2,3-Butanediol (BD) |
| 2,3-Butanediol (BD) | 2-Methoxyethanol (ME) |
| 2-Methoxyethanol (ME) | Acetamide (AM) |
| 2-Methyl-1,3-Propanediol (MP) | Diethylene Glycol (DG) |
| 2-Methyl-2,4-Pentanediol (MPD) | Dimethyl Sulfoxide (DMSO) |
| Acetamide (AM)* | Ethylene Glycol (EG) |
| Diethylene Glycol (DG) | Formamide (FA) |
| Diglyme | Glycerol (Gly) |
| Dimethyl Sulfoxide (DMSO) | N-Methylacetamide (NMA) |
| Dimethylacetamide (DMA) | Propylene Glycol (PG) |
| Ethylene Glycol (EG) | Binary CPA mixtures composed of the 11 CPAs above (equimolar split between CPAs). We tested 46 out of 55 possible binary mixtures. We omitted the following binary mixtures because they were tested in our previous study [35]: DMSO/EG, DMSO/FA, DMSO/Gly, DMSO/PG, EG/FA, EG/Gly, EG/PG, FA/PG, Gly/PG. |
| Formamide (FA) |  |
| Glycerol (Gly) |  |
| N-Methylacetamide (NMA) |  |
| Propionamide (PA) |  |
| Propylene Glycol (PG) |  |
| Tetraethylene Glycol Dimethyl Ether (TGDE) |  |
| Tetrahydrofurfuryl Alcohol (TFA) |  |
| Triethylene Glycol (TG) |  |
| Triglyme |  |

\* Acetamide had a 1.1 mol/kg concentration because of an error during solution preparation.

### Experimental Overview

This study builds on the methods outlined in our previous study [35], utilizing a Hamilton Microlab STARlet automated liquid handling system for CPA addition and removal. The Hamilton system was used to ensure precise, automated solution exchanges, with experimental conditions adapted to evaluate the 21 selected CPAs and their binary mixtures. As in our previous study, experiments were performed on bovine pulmonary artery endothelial cells (BPAEC) cultured in 96 well plates. To evaluate CPA toxicity, viability was first assessed prior to CPA exposure using the PrestoBlue assay, a resazurin-based viability assay where metabolically active cells reduce nonfluorescent resazurin into fluorescent resorufin. The cells were then exposed to CPA using the automated liquid handling system, and a final PrestoBlue measurement was made 24 h after CPA exposure. Cell viability was calculated using the double normalization process described in our previous study: the final PrestoBlue reading was normalized to the initial PrestoBlue reading from the same well, as well as the mean PrestoBlue reading for the CPA-free control wells. Compared to our previous study, which tested 15 CPA compositions per plate, the well plate layout in the current study was adjusted to accommodate a greater number of CPA compositions. Each plate included 19 CPA compositions (4 replicate wells each), along with wells designated for a CPA-free positive control (8 wells), negative control (4 wells), and cell-free background control (8 wells). To streamline the experimental process, the CPA addition and removal protocols were modified and simplified compared to our previous study [35]. CPA addition and removal for a 3-mol/kg concentration consisted of direct exposure to 3 mol/kg CPA in isotonic buffer, followed by exposure to hypertonic buffer (1,200 mOsm by adding extra NaCl) for 4.5 min before returning to isotonic conditions. For comparison, we also assessed our previous multistep method [35], as well as single step exposure to 3 mol/kg CPA. CPA addition and removal for 6 mol/kg CPA consisted of exposure to 3 mol/kg CPA in isotonic buffer for 2.5 min, 6 mol/kg CPA in isotonic buffer for 30 min, 3 mol/kg CPA in hypertonic buffer for 8 min, and finally to the hypertonic buffer for 10 min, before returning to isotonic conditions. Figure 1 illustrates these CPA addition and removal methods.

**Figure 1.**
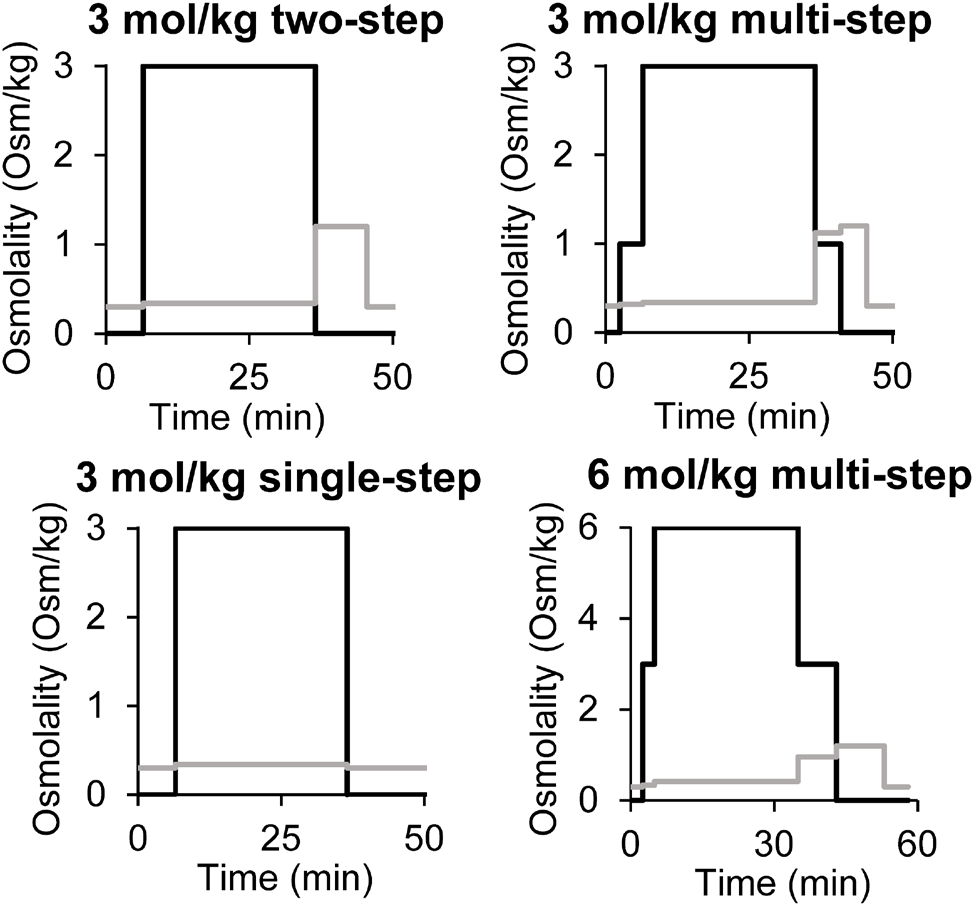
CPA addition and removal methods. CPA concentration (mol/kg) is shown with the black lines and nonpermeating solute concentration (osmol/kg) is shown with the gray lines.

All other methods were carried out as described in our previous study, with one exception: the pH of the base HEPES-buffered solutions used for the CPA preparations was not adjusted to 7.3 due to an error. The actual pH of these solutions was estimated to be ∼5.5 afterwards by preparing the buffer in the same way and measuring the resulting pH. Despite the pH difference, we obtained similar viability results in the current study and previous pH-adjusted study for the isotonic control, indicating that exposure to the relatively low pH did not compromise cell viability. In particular, we obtained a final PrestoBlue reading that was 1.9 times higher than the initial PrestoBlue reading, which shows that the cells in the isotonic control wells approximately doubled during the 24 h culture period.

### Statistical Analysis

Statistical analyses were performed using the Python packages statsmodels and SciPy. As in our previous study [35], we used the boxplot method for outlier detection. We used the standard interquartile range coefficient of 1.5 to identify outliers. Data are presented as mean ± standard error of the mean (SEM), as indicated. For most comparisons involving multiple groups, one-way analysis of variance (ANOVA) was used, followed by Tukey’s post hoc tests for pairwise comparisons. A p-value below 0.05 was considered statistically significant. For analysis of toxicity reduction in CPA mixtures, a different approach was used due to the large number of comparisons. In this case, Welch’s two-sample, two-sided t-test was performed, followed by the Benjamini– Hochberg method as a correction for multiple testing [6]. A corrected p-value (q-value) < 0.05 was considered statistically significant. The normality assumption for the data was assessed using Q-Q plots. Visual inspection of the Q-Q plots indicated that the normality assumption was valid for all groups analyzed.

## 3. Results and Discussion

We recently presented a new high throughput method for simultaneous measurement of CPA permeability and CPA toxicity [1]. We used this method to screen 27 chemicals, resulting in 21 with favorable permeability and toxicity properties after exposure to 2 mol/kg CPA at room temperature. In the current study, we examine the toxicity of these 21 chemicals at higher concentrations, and we explore the potential for combining the CPAs in mixtures to reduce toxicity. For these experiments, we used a high throughput toxicity assay that leverages robotic liquid handling to automate the CPA addition and removal process [35].

We initially examined whether the experimental workflow for adding and removing CPA could be simplified to facilitate scalability in future screening processes. To address this, we compared three techniques for exposure to 3 mol/kg CPA: a single-step method, a two-step method, and the multi-step method that we used in our previous study [35]. Figure 2 displays the experimental comparisons among these three approaches for a selection of compounds. Out of all the CPAs that were examined, glycerol has the lowest membrane permeability [1,17,20,30,31,34,35], which results in the most notable alterations in volume and the highest risk of osmotic damage. The viability of cells exposed to 3 mol/kg glycerol for 30 minutes was significantly higher for both multi-step and two-step processes compared to the single-step process, which is consistent with osmotic damage in the latter case. However, the cell viability values for the two-step and multi-step methods were not statistically different, suggesting that the two-step method is sufficient to prevent osmotic damage. The cell viability after exposure to DMSO, PG, and EG was close to 100% in all cases, suggesting negligible osmotic damage for these CPAs. For FA, there was significant loss of viability for all the CPA addition and removal methods, and the multistep method was the most damaging. This is consistent with increased toxicity due to prolonged CPA exposure during the multistep method. Overall, the results demonstrate that the two-step process is superior to the alternative methods. As a result, we opted to use the two-step method in subsequent experiments.

**Figure 2.**
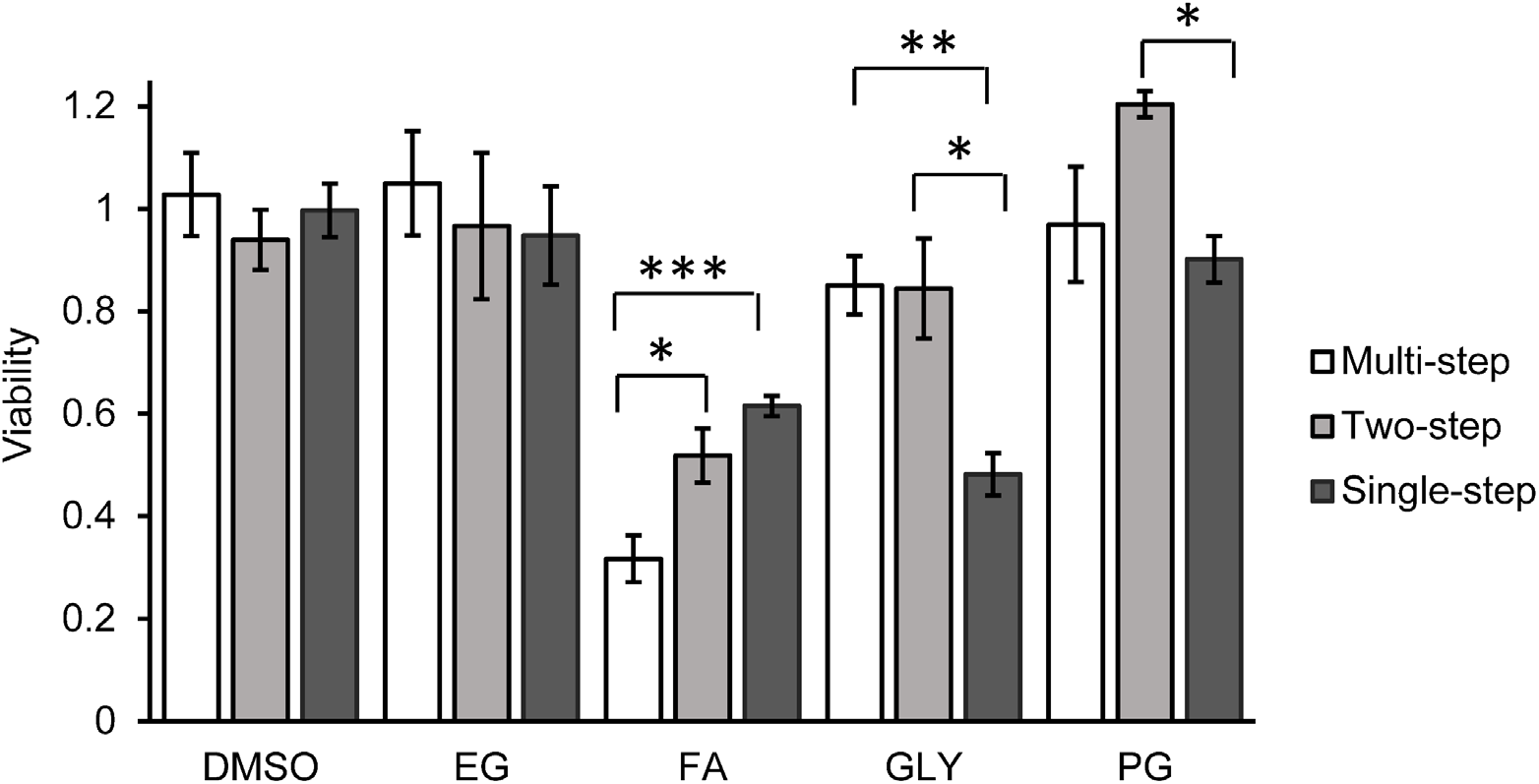
Comparison of CPA addition and removal methods for exposure to 3 mol/kg CPA. Each bar represents the mean ± SEM from 3-5 replicate wells. Experiments were conducted at room temperature with a 30-minute exposure time. Statistically significant differences between groups are indicated by asterisks, with the following p-value thresholds: * p < 0.05, ** p < 0.01, *** p < 0.001.

Figure 3 illustrates the cell viability for a CPA concentration of 3 mol/kg, with two distinct exposure times of 10 and 30 minutes. Of the 21 CPAs tested, only 7 yielded a viability exceeding 80% after 30 min exposure (GLY, DMSO, PG, EG, DG, PD, and ME). However, cell viability was higher when the CPA exposure time was reduced to 10 min. Overall, the toxicity data for 3 mol/kg CPA was used to select 11 CPAs for subsequent testing of single-CPA solutions and binary CPA mixtures at a concentration of 6 mol/kg. Due to the large number of conditions, these experiments were limited to a single exposure time of 30 min.

**Figure 3.**
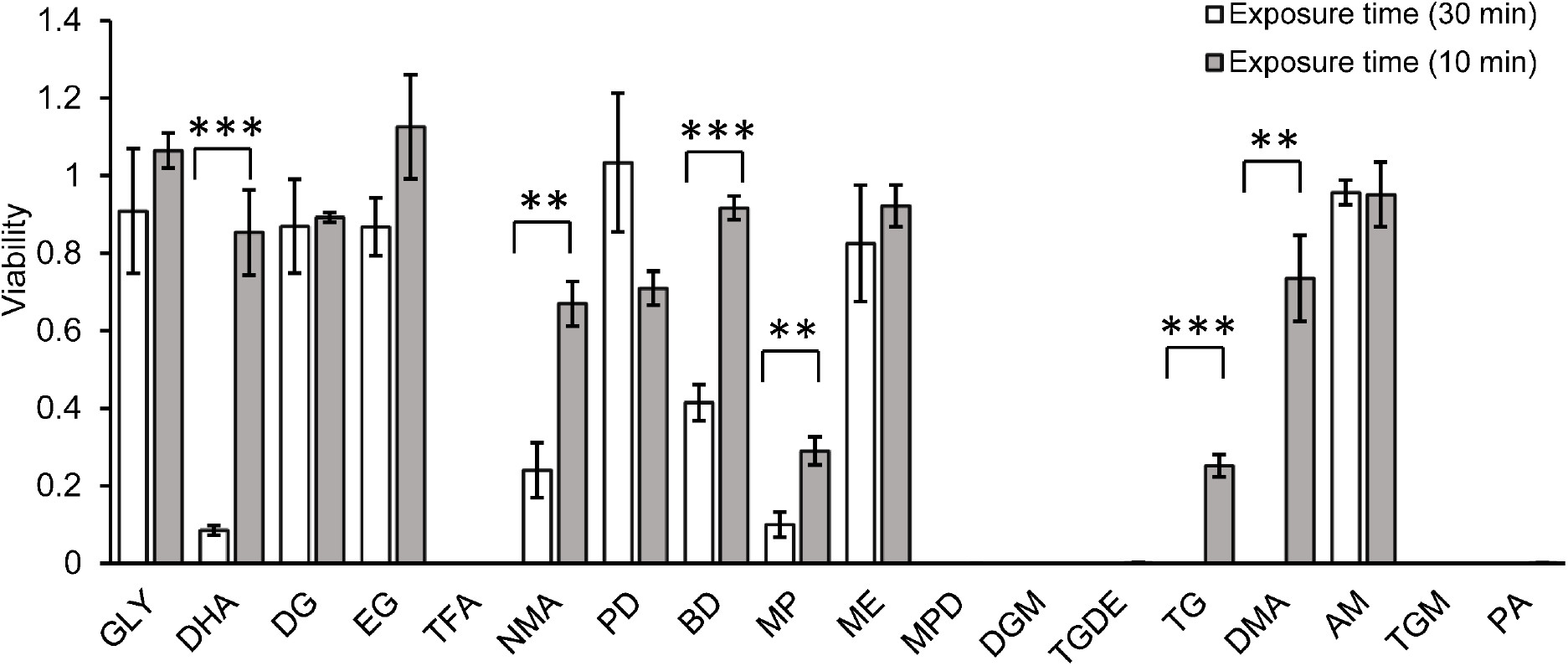
Effect of exposure time on cell viability after exposure to 3 mol/kg CPA. Each bar represents the mean ± SEM from 3-4 replicate wells. Experiments were conducted at room temperature. Statistically significant differences between groups are indicated by asterisks, with the following p-value thresholds: * p < 0.05, ** p < 0.01, *** p < 0.001. Note: The concentration of acetamide was 1.1 mol/kg due to an error.

Figure 4 compares the toxicity of 3 mol/kg and 6 mol/kg solutions containing a single CPA. Of the 11 CPAs tested, only EG yielded high viability after 30 min exposure to a 6 mol/kg concentration. For the other CPAs, the cell viability was significantly lower after exposure to 6 mol/kg CPA compared to the viability after exposure to 3 mol/kg CPA. This highlights the concentration-dependence of CPA toxicity.

**Figure 4.**
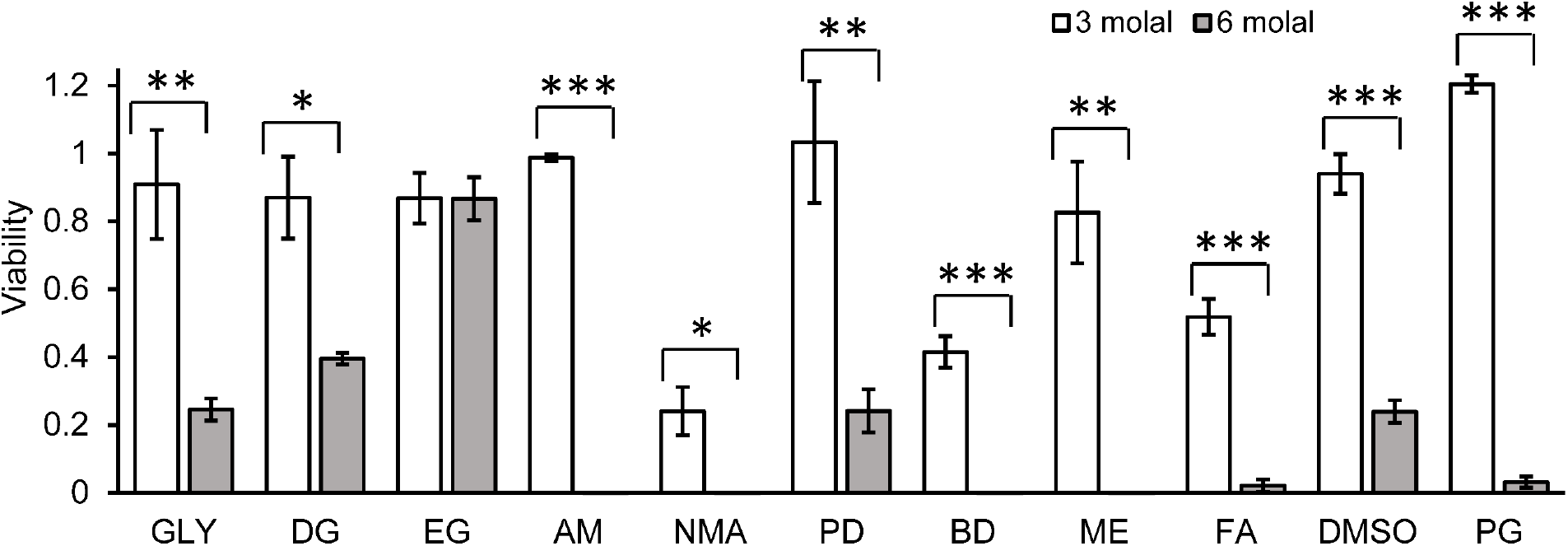
Effect of CPA concentration on cell viability after 30 min CPA exposure. Each bar indicates the mean ± SEM from 3-4 replicate wells. Experiments were conducted at room temperature. Statistically significant differences between groups are indicated by asterisks, with the following p-value thresholds: * p < 0.05, ** p < 0.01, *** p < 0.001. Note: For acetamide, the bar for 3 mol/kg actually represents the viability after exposure to a 1.1 mol/kg concentration due to an error during solution preparation.

Previous studies have suggested that CPA mixtures may exhibit reduced toxicity compared to equivalent concentrations of their constituent CPAs [2,3,12,14,15,24,33,35]. To explore the potential for toxicity reduction in mixtures, we compared the toxicity of individual CPAs with their corresponding binary mixtures, both at a concentration of 6 mol/kg. Overall, we assessed 46 CPA mixtures, as shown in Figure 5. For 20 of the 46 mixtures, cell viability was higher after exposure to the mixture than both corresponding single CPA solutions. This suggests possible toxicity-reducing effects for these CPA combinations. In contrast, only 2 of the CPA mixtures exhibited lower viability than the equivalent concentration of both corresponding single CPA solutions, and in these cases the viability difference was less than 4%.

**Figure 5.**
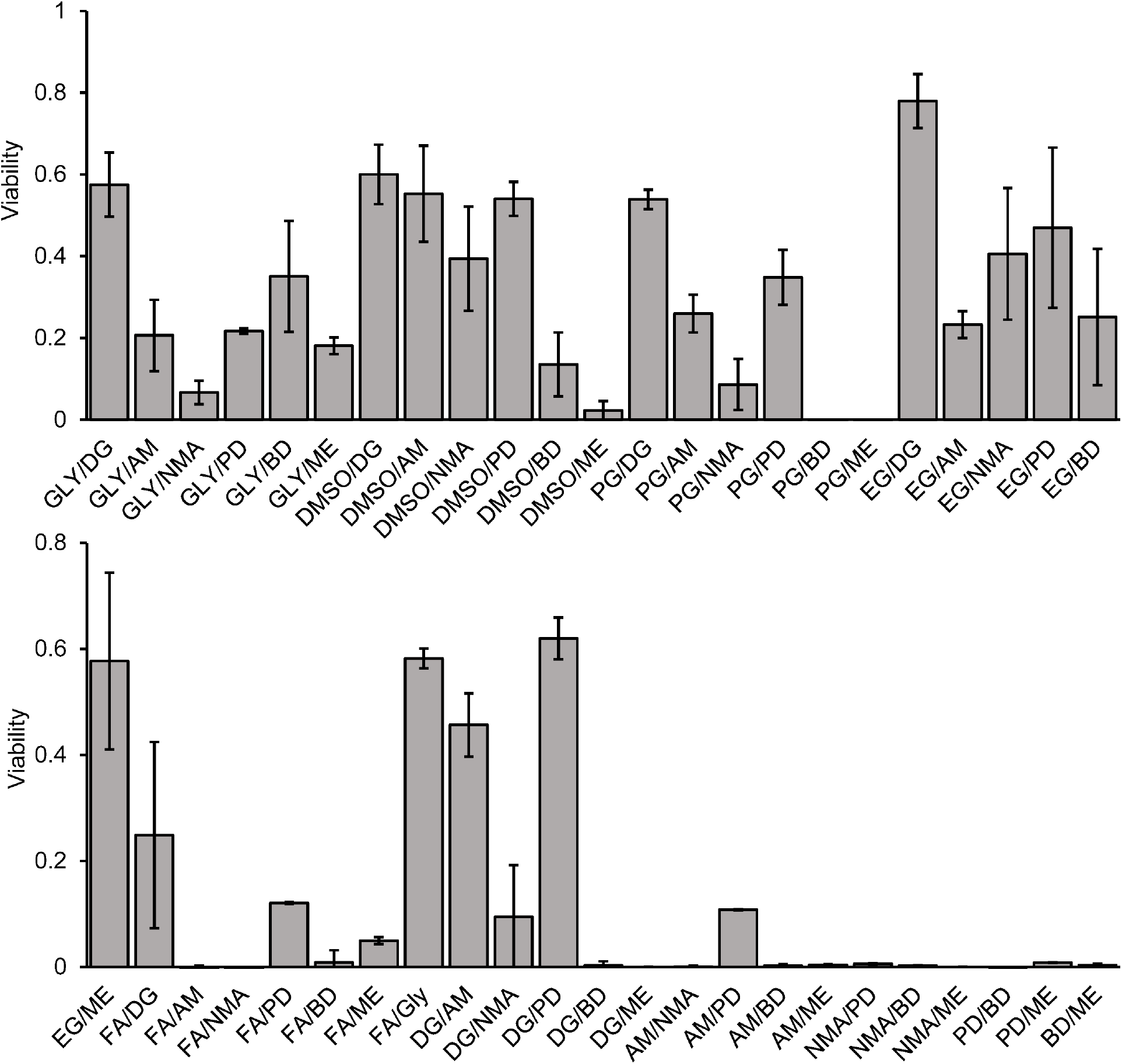
Cell viability after exposure to 6 mol/kg CPA mixtures for 30 min at room temperature. Each bar indicates the mean ± SEM from 3-4 replicate wells.

To quantitatively examine toxicity reduction in mixtures, we statistically compared mixtures containing two CPAs at a total concentration of 6 mol/kg to both the corresponding single-CPA solutions at a concentration of 6 mol/kg. Overall, we did over 100 t-tests to compare each of the 46 mixtures to the corresponding single CPA solutions. To limit the proportion of false positive results to less than 5%, we used false discovery rate analysis as described by Benjamini and Hochberg [6]. This analysis revealed eight cases where the viability for the binary CPA mixture was statistically higher than the viability for both individual CPAs. Four of these cases are illustrated in Figure 6. In the remaining four cases, including AM/ME, BD/ME, NMA/BD, and NMA/ME, the maximum viability was less than 2 percent. Overall, this comparison between single-CPA solutions and binary CPA mixtures reveals that, in many cases, mixtures reduce toxicity.

**Figure 6.**
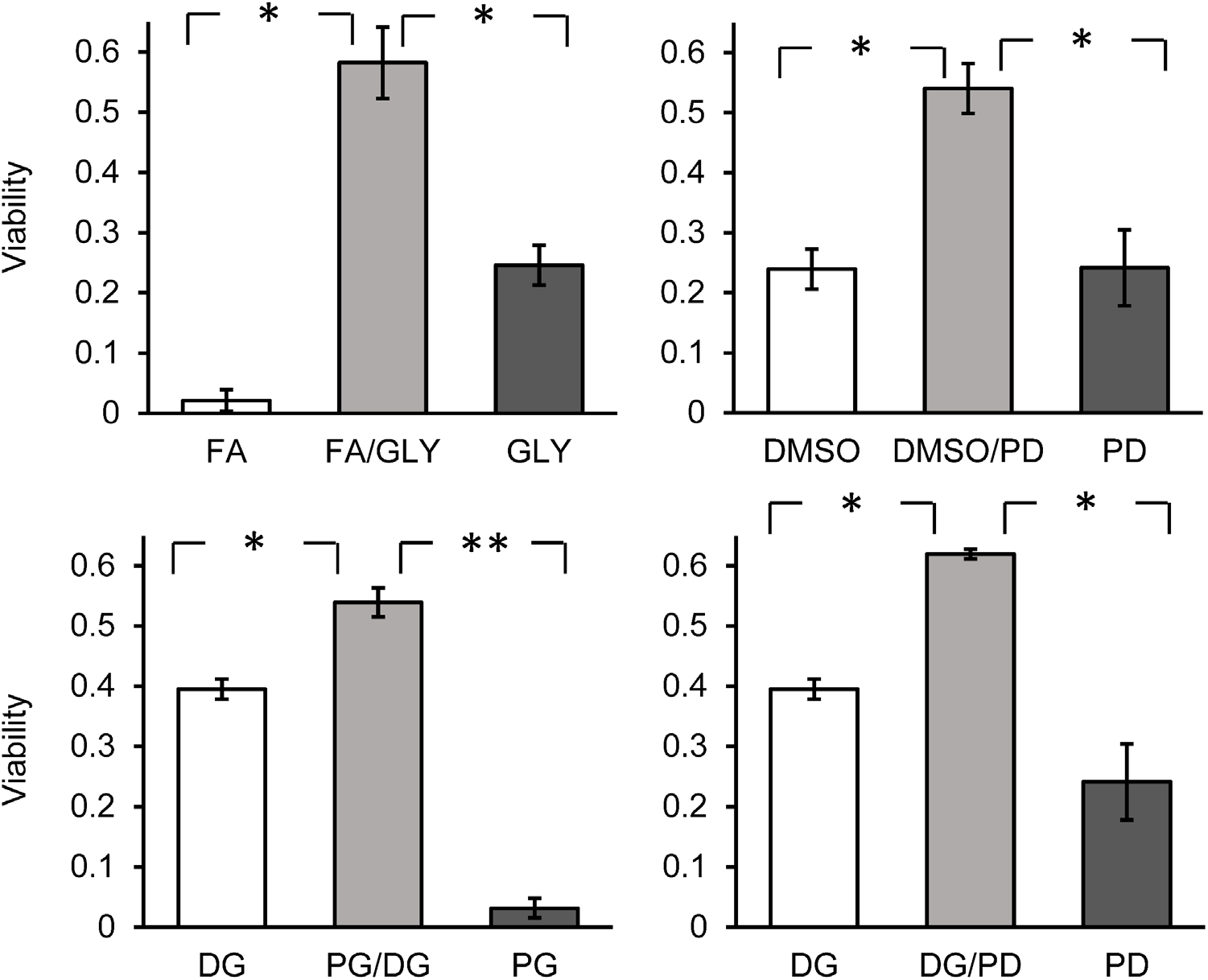
Viability for 6 mol/kg CPA mixtures compared to their 6 mol/kg individual CPA constituents. Each bar indicates the mean ± SEM from 3-4 replicate wells. Experiments were conducted at room temperature. Statistically significant differences between groups are indicated by asterisks, with the following p-value thresholds: * p < 0.05, ** p < 0.01.

Previous studies have reported toxicity neutralization in some cases, including the neutralization of FA toxicity by addition of either DMSO or GLY [12,35]. Toxicity neutralization occurs when the addition of a second CPA reduces or eliminates the toxicity of the first CPA. In the current study, toxicity neutralization was evaluated by comparing the viability of cells exposed to a 3 mol/kg single-CPA solution with the viability of cells exposed to a 6 mol/kg binary CPA solution containing 3 mol/kg of each CPA. This analysis did not reveal any cases of statistically significant toxicity neutralization. However, there were three cases where the 6 mol/kg mixture yielded higher viability than one of the 3 mol/kg single-CPA solutions, including DMSO/NMA (p = 0.35), EG/NMA (p = 0.40), and FA/GLY (p = 0.45). While our previous study [35] reported toxicity neutralization at higher concentrations for FA/GLY, further investigation is needed to confirm this observation. It is important to note that DMSO/FA, a combination previously shown to exhibit toxicity neutralization, was not tested in this study.

## 4. Conclusions

Our investigation underscores the potential of high-throughput screening of CPA mixtures in advancing vitrification techniques for cryopreserving complex biological structures. The study highlights that while traditional vitrification methods rely on a limited range of CPAs, exploring a broader array of compounds has the potential to yield improvements. Specifically, our findings demonstrate that CPA toxicity increases with both concentration and exposure duration, and that CPA mixtures reduce toxicity in many cases. This suggests that carefully selected combinations of CPAs can significantly improve vitrification outcomes and reduce associated toxicity. Our findings align with several established patterns, such as the relative non-toxicity of ethylene glycol, the higher toxicity of formamide, and the observation that several CPA mixtures diminish overall toxicity.

Building on these findings, future research should focus on broadening the chemical library of potential CPAs by testing a wider range of novel compounds. Future efforts should also examine the potential for reducing CPA toxicity by lowering the temperature. While the current study examined CPA toxicity at room temperature, the Hamilton system can be modified to enable toxicity measurement at subambient temperatures. Overall, high-throughput screening of CPA toxicity has the potential to generate large datasets that can serve as foundation for future efforts to develop predictive models for more efficient identification and evaluation of new CPAs. These advancements have the potential to improve vitrification protocols, enhancing the preservation of complex biological structures and expanding their applications in both medical and research fields.

## Supporting information

Viability data

## 5. Acknowledgements

This work was supported by funding from the National Institutes of Health (R01 EB027203) and by a donation from the Cryonics Institute to the laboratory of AZH.

## Notes

### Competing Interest Statement

The authors have declared no competing interest.

